## Supplemental Material for "Relative Distribution of DnaA and DNA in *Escherichia coli* Cells as a Factor of Their Phenotypic Variability" (from another version of this preprint: version 2, 2024-09-23)

### Construction of MG1655 (*dnaA-egfp*, *hu-mcherry*).

*Insertion of kanamycin resistance selectable marker.* The *kanR* gene was amplified by PCR with two primers, 5'ataattaGCTAGCGAGATCCGGCTGCTAACA3' and 5'attataaGCTAGCCAGAGTTCTTGAAGTGGTGG3', using pET28a as a template. The PCR product and pBAD24-dnaA-mCherry plasmid (described in (1)) were cut with NheI and subsequently ligated. This placed *kanR* before *dnaA-mCherry* gene for the construction of pBAD24-kanR-dnaA-mCherry plasmid.

*Insertion of DnaA promotor*. The promotor was amplified by PCR with two primers, 5'attataaGCTAGCGTAAAGCGCAAGGATCGTC3' and 5'CCCGTACCCAATCGAGGACAA3', using MG1655 chromosome as a template. pBAD24-kanR-dnaA-mCherry plasmid and the PCR product were cut with NheI and EcoRI and subsequently ligated, resulting in pBAD24-kanR-dnaA-promotor-dnaA-mCherry plasmid.

*Replacement of mCherry with egfp*. *egfp* was amplified by PCR with two primers, 5'taatatGGTACCATGGTGAGCAAGGGCGAGGAG3' and 5'attataCTCGAGCTTGTACAGCTCGTCCATGCC3', using pEGFP vector as a template (Clontech Laboratories, Inc., Palo Alto, CA). pBAD24-kanR-dnaA promotor-dnaA-mCherry and the PCR product were cut with KpnI and XhoI and subsequently ligated, resulting in pBAD24-kanR-dnaA promotor-dnaA-egfp plasmid.

*Chromosomal insertion of kanR-dnaA promotor-dnaA-egfp cassette.* The cassette was amplified by PCR using 5'ATGGGTAAAGCGCAAGGATCGTCCTGGATCTTTATTAGATCGATTAAGCCGGCGTCACACTTTGCTAT3' and 5'GCACCGCCAGGGTTATCCG3' primers. The resulting PCR product was inserted into *dnaA* locus of wtMG1655 by the Recombineering technique as described in (2), resulting in MG1655(*dnaA-egfp*). The chromosomal insertion was verified by PCR and fluorescence microscopy imaging.

*Chromosomal insertion of mCherry-ampR cassette*. The cassette was amplified by PCR with two primers, 5’GTACCGGCATTTGTTTCTGGCAAGGCACTGAAAGACGCAGTTAAGtctcgcATGGTGAGCAAGGGCGAGGAG3’ and 5’CCAGCCAGCATCAATGATCGACGCCAGAAAGACAAAAGGGGTGAAACCACCTGGCAAGTGTAGCGGTCACG3’, using pBAD24-mCherry as a template (described in (1)). The resulting PCR product was inserted in-frame downstream of *hupA gene* of MG1655 (*dnaA-egfp*) by Recombineering technique as described (2) thus producing MG1655 (*dnaA-egfp*, *hu-mcherry*). The chromosomal insertion was verified by PCR and fluorescence microscopy imaging for the expression of both fluorescent proteins.

The functionality of both DnaA-eGFP and HU-mCherry fusions was confirmed by comparison of the strains' growth rates and size distributions with the wtMG1655 (Fig. S1).

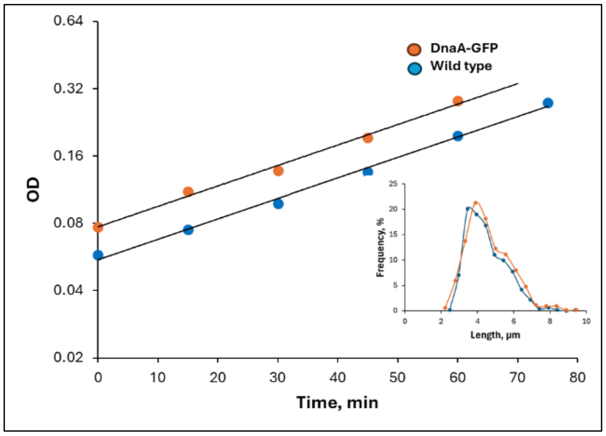


**Fig. S1.** Growth curves of MG1655 (*hu-egfp*) (blue dots) and double-tagged MG1655 (*dnaA-egfp*, *hu-mcherry*) (orange dots) in Neihardt’s full medium (basal plus supplement) show the same generation time of 32 min as wtMG1655 (not shown). Size distributions of both strains (inset) coincide well with each other.

### Quantitative characteristics of pixel-wise colocalization.

The Pearson coefficient shows the strength and mode of the correlation, while the Manders coefficient shows the degree of overlap between the two species (3).

**Table S1.** Pearson’s and Mander’s coefficients for cells of different sizes (or ages) at various functional states. The coefficients were generated by the ImageJ plugin JaCoP (4) (https://imagej.net/plugins/jacop)

| Type of treatment | Type of cell | Pearson coefficient | Manders coefficient (fraction of HU overlapping DnaA) | Manders coefficient (fraction of DnaA overlapping HU) |
|---|---|---|---|---|
| **Untreated cell** | Short | 0.889 | 0.984 | 0.7 |
| | Medium | 0.808 | 0.979 | 0.638 |
| | Long | 0.856 | 0.986 | 0.765 |
| **CAM-treated cell** | Short | 0.653 | 0.997 | 0.449 |
| | Medium | 0.646 | 0.995 | 0.496 |
| | Long | 0.653 | 0.997 | 0.449 |
| **NAL-treated cell** | Short | 0.764 | 0.972 | 0.584 |
| | Medium | 0.692 | 0.942 | 0.556 |
| | Long | 0.655 | 0.916 | 0.354 |
| **NAL+CAM cell** | Short | 0.318 | 0.853 | 0.685 |
| | Medium | 0.48 | 0.991 | 0.277 |
| | Long | 0.14 | 0.933 | 0.139 |
| **RIF-treated cell** | Short | 0.895 | 0.958 | 0.858 |
| | Medium | 0.966 | 0.996 | 0.899 |
| | Long | 0.955 | 0.986 | 0.884 |

### Intracellular asymmetry in DNA distribution in wtDnaA strain

The asymmetry indexes of DNA distribution in cells of the MG1655 (*hu-egfp*) strain containing the wtDnaA were calculated as those shown in Table 2 of the main text, except that GFP fluorescence was used to measure the HU distribution.

**Table S2.** Population averages (± SD) of concentration asymmetry in MG1655 (*hu-egfp*).

| Leader pole | Asymmetry index | | |
|---|---|---|---|
| | Volume | HU | DAPI |
| **Diameter** | 1.15 ± 0.10 | 0.95 ± 0.19 | 0.97 ± 0.32 |
| **HU** | 1.07 ± 0.17 | 1.10 ± 0.22 | 1.11 ± 0.35 |
| **DAPI** | 1.07 ± 0.17 | 1.08 ± 0.22 | 1.13 ± 0.35 |
| **None** | 1.05 ± 0.17 | 1.00 ± 0.20 | 1.02 ± 0.33 |

### Examples of FRAP measurements of the free eGFP, DnaA (L417P), and DnaA and HU in CAM- or Rif-treated cells

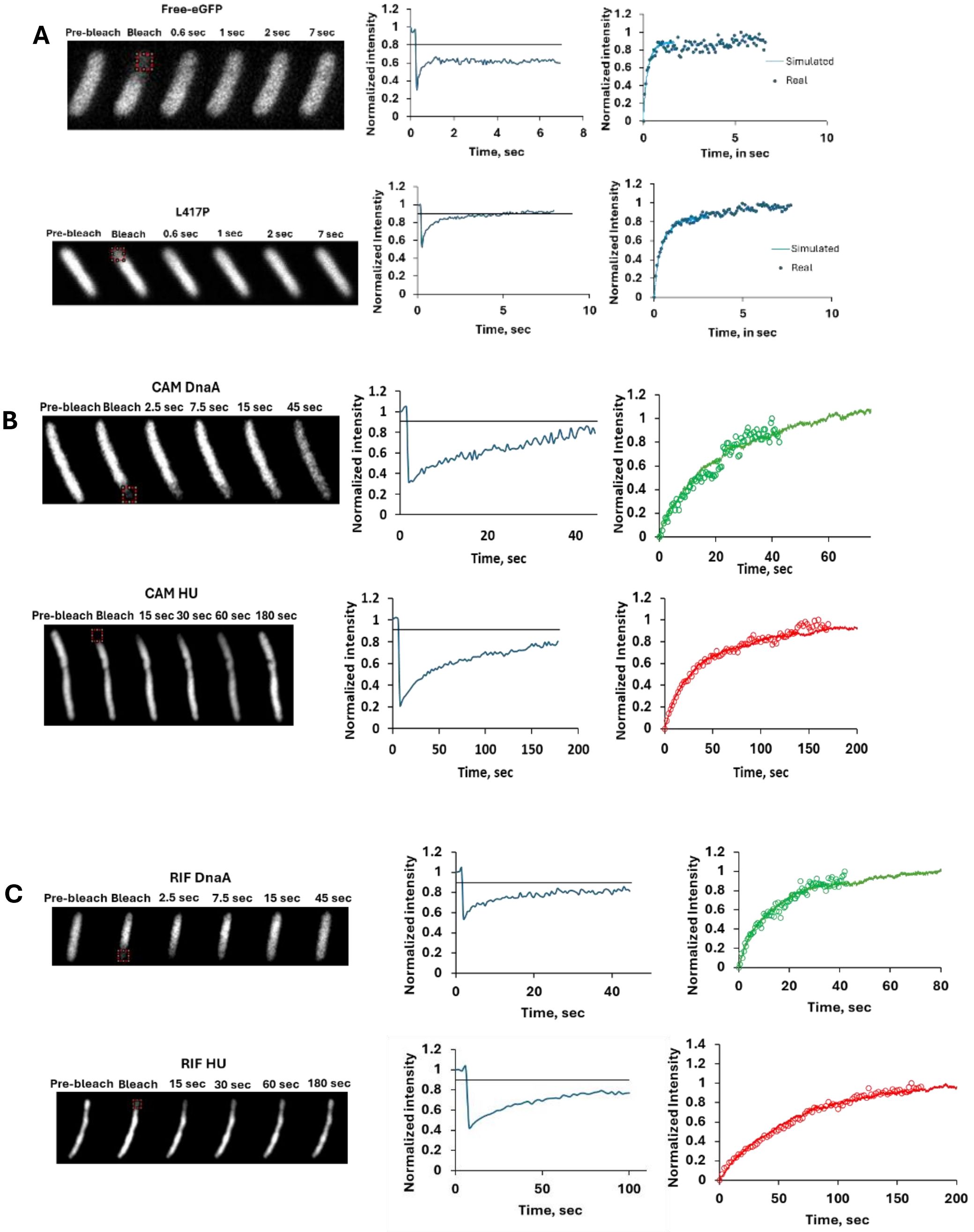

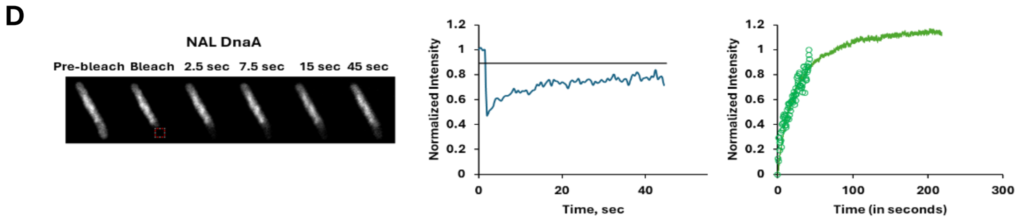


**Fig S2.** (**A**) Free-eGFP and mutant DnaA (L417P) cells, (**B**) CAM-treated cells, (**C**) RIF-treated cells, and (**D**) NAL-treated cells. The cells were bleached on poles (1x1 µm red squares on the second image in the series) and the fluorescence recovery in these areas (blue lines in the central column) was further analyzed by simFRAP ((5) and see Methods, Section 4) (plots in the right column: green circles and line – real and simulated DnaA recovery kinetics, red – the corresponding curves for HU). The horizontal lines in recovery plots show the expected recovery maxima.

### Cell-to-cell variation of DnaA and HU in slow-growing or antibiotic-treated cultures.

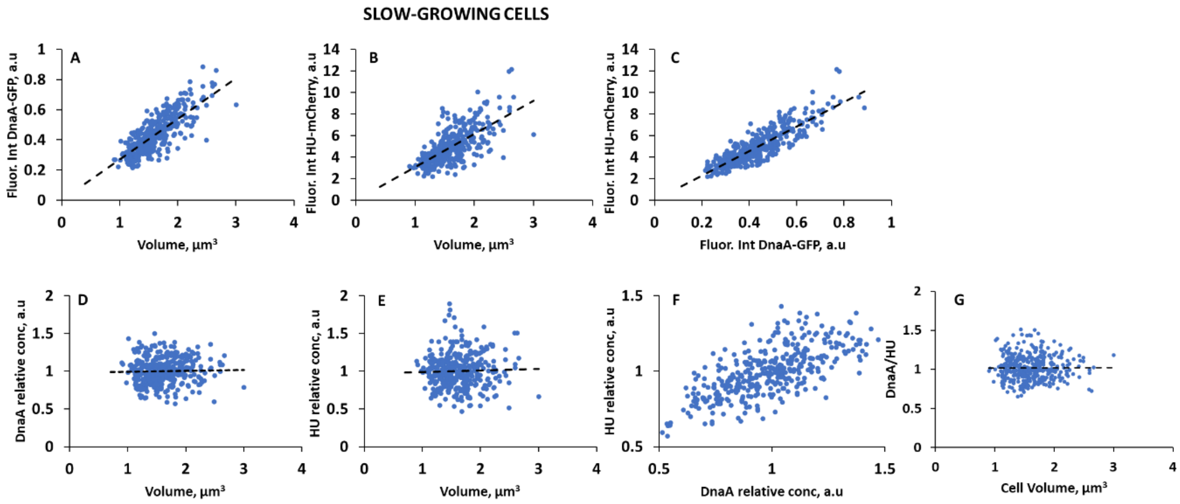


**Fig S3.** DnaA and HU amounts and concentrations in cells in a slow-growing population. Approximately 500 cells from a steady-state culture grown in the Neidhardt medium without supplement (generation time 65 min) were measured. The image analysis was performed using ImageJ with the ObjectJ plugin and Coli-inspector protocol ((6) and see Methods for details). The dependences of total cellular fluorescence intensities of DnaA-GFP **(A)** and HU-mCherry **(B)** on cell volume and their mutual relation **(C)** are shown. The data were fitted to a linear function. The corresponding dependences of DnaA **(D)** and HU **(E)** concentrations (normalized to the population average) on cell volume are shown. **F** - The relationship between DnaA and HU normalized concentrations. **G** – Cell volume dependence of DnaA/HU.

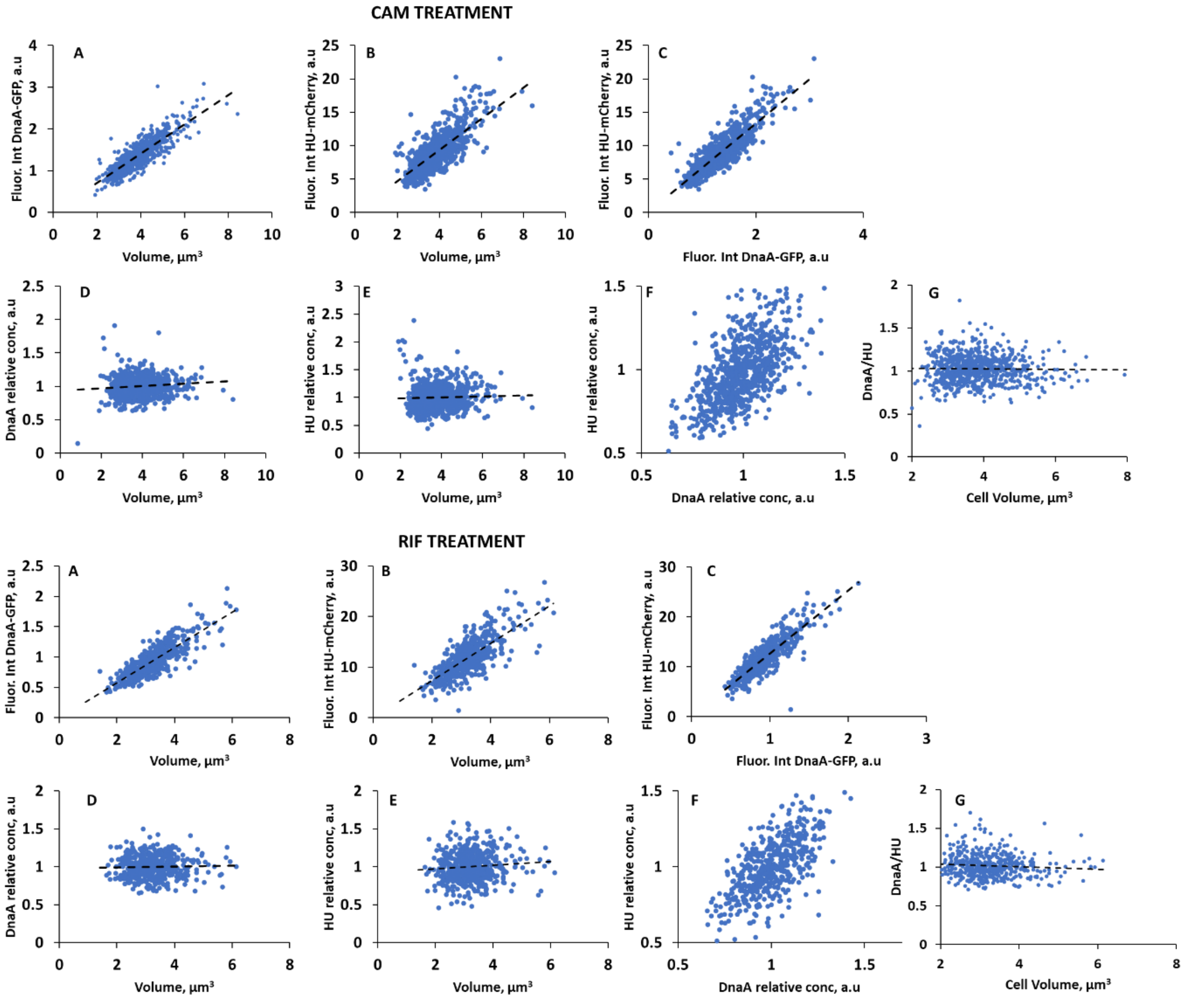


**Fig S4.** DnaA and HU amounts and concentrations in cells in CAM- and RIF-treated populations. Portions of steady-state culture grown in the full Neidhardt medium (generation time 32 min) were treated with CAM or Rif (see Methods). The image analysis was performed using ImageJ with the ObjectJ plugin and Coli-inspector protocol ((6) and see Methods for details). The dependences of total cellular fluorescence intensities of DnaA-GFP **(A)** and HU-mCherry **(B)** on cell volume and their mutual relation **(C)** are shown. The data were fitted to a linear function. The corresponding dependences of DnaA **(D)** and HU **(E)** concentrations (normalized to the population average) on cell volume are shown. **F** - The relationship between DnaA and HU normalized concentrations. **G** – Cell volume dependence of DnaA/HU.

**Table S3. The variance of normalized concentrations of DnaA and HU and their ratio in CAM- and RIF-treated populations**

| | | **CAM** (CV) | **RIF** (CV) |
|---|---|---|---|
| **HU** | | 0.037 (0.21) | 0.036 (0.19) |
| **DnaA** | | 0.020 (0.14) | 0.020 (0.14) |
| **DnaA/HU** | population | 0.022 (0.15) | 0.025 (0.16) |
| | young* | 0.028 (0.17) | 0.026 (0.16) |
| | adult* | 0.017 (0.13) | 0.025 (0.16) |

*Young – subpopulation of cells smaller than median volume; Adult - larger than median volume.

### Distribution of DNA material in a CAM-treated cell

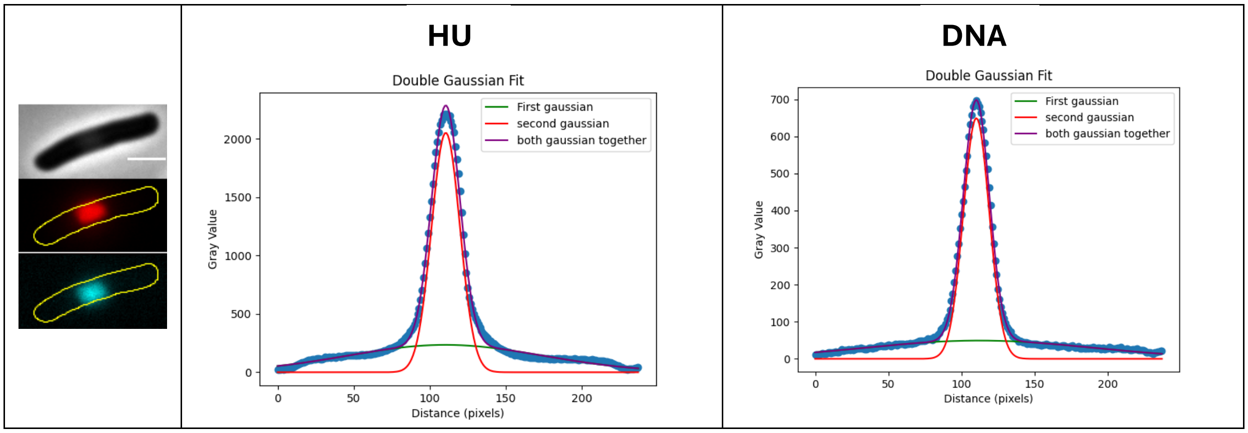


**Fig S5.** Deconvolution of fluorescence intensity profiles of HU and DAPI in a CAM-treated cell (left column). The profiles were measured along the cell image (red – HU fluorescence, cyan – DAPI, false colors) with a strip 5 pixels wider than the cell width. The profile values (blue dots) were fitted with two Gaussians using Python program. The Gaussian parameters are obtained as follows. For HU: amplitude1=234.6 mean1=110.3 std dev1=64.1, amplitude2= 2052.1, mean2=110.6, std dev2=9.3. For DNA: amplitude1=49.3, mean1=112.1, std dev1=77.7, amplitude2=649.2, mean2=110.1, std dev2= 8.9.
